# Induction of promoter methylation and transcriptional gene silencing upon high pressure spraying of 24-nt small RNAs in *Nicotiana benthamiana*

**DOI:** 10.1101/2022.03.14.484340

**Authors:** Melanie Walz, Linus Hohenwarter, Gabi Krczal, Michael Wassenegger, Veli Vural Uslu

## Abstract

Exogenous RNA application is a promising transgene-free approach for crop protection and improvement. In almost all applications reported so far, exogenous RNA molecules (double stranded RNAs or small RNAs) are applied to plants in order to trigger degradation of a given mRNA (of plant, pest or pathogen origin), in a process termed post-transcriptional gene silencing (PTGS). However, whether exogenous RNAs can also trigger epigenetic modifications to plants such as RNA-directed DNA methylation (RdDM) and transcriptional gene silencing (TGS) remains largely unaddressed. Here, we provide evidence that high pressure spraying of a 24-nt short interfering RNA (siRNA) designed to target the CaMV 35S promoter that drives the expression of a GFP transgene, resulted in promoter RdDM in *N. benthamiana*. Moreover, the methylation at the target site spreads both up and downstream neighboring sites. Importantly, GFP expression was reduced in the sprayed leaf tissues. Small RNA sequencing excluded the presence siRNAs mapping to the GFP gene body and documented substantial amount of secondary RNAs in the promoter, suggesting that the GFP silencing was transcriptional rather than post-transcriptional. Our study provides a proof of principle for <u>sp</u>ray <u>i</u>nduced <u>e</u>pigenetic <u>m</u>odification (SPIEM) that could be used in modern crop breeding platforms.

## INTRODUCTION

RNA interference (RNAi) is a fundamental process in plants for development and stress response. Plants use small RNAs (sRNAs) for post-transcriptional gene silencing (PTGS) by cleaving the target transcripts or by inhibiting their translation into proteins (Brodersen *et al*., 2008; Hamilton and Baulcombe, 1999; Hung and Slotkin, 2021). Besides, target transcripts can also be silenced even before they are produced by transcriptional gene silencing (TGS) via epigenetic modifications specifically at their promoter regions based on sequence complementarity (Matzke *et al*., 2015; Mette *et al*., 2000; Wassenegger, 2005) (Vaucheret and Fagard, 2001).

Two main classes of sRNAs undertake RNAi: Micro RNAs (miRNAs) and siRNAs. MiRNAs are transcribed as pri-miRNAs by RNA polymerase 2 (Pol II). Pri-miRNAs fold back on themself forming a hairpin-like structure, which are further processed by mostly Dicer-like endonuclease 1 (DCL1) into short duplex RNA structure with 2nt overhangs (Wang et al 2019). On the other hand, siRNAs are mostly produced through processing of dsRNA, which may originate from inverted repeats or viral replication intermediates. DCL4, DCL2, and DCL3 cleave dsRNAs into 21-nt, 22-nt, and 24-nt small RNAs (sRNAs), respectively (Henderson et al 2006, Sanan-Mishra et al 2021). Most miRNAs and 21-22nt long siRNAs are loaded onto ARGONAUTE proteins and assemble RNA-induced silencing complex (RISC) for PTGS (Baulcombe 2004, Voinnet, 2009). RISC executes direct cleavage of target transcripts and/or recruit RNA-directed-RNA polymerase (RDR) for re-amplification of double stranded RNA. Newly produced dsRNA is again cleaved into secondary sRNAs, which extend beyond the initial target site, which is known as transitivity. Therefore, the positive feedback loop assures efficient and rapid removal of the target transcript via PTGS (de Felippes and Waterhouse 2020).

In plants, PTGS is tightly connected to RdDM and TGS (Baulcombe, 2004; Jones *et al*., 1999). RdDM is triggered primarily by 24-nt siRNAs, although other siRNA classes and/or larger dsRNA molecules may also be involved in the process (Dalakouras and Vlachostergios, 2021; Gallego-Bartolome, 2020; Matzke *et al*., 2015; Sigman *et al*., 2021; Wassenegger and Dalakouras, 2021). According to the most recent model, AGO4-loaded 24-nt siRNAs are suggested to recruit RNA Polymerase V (Pol V) to the target DNA locus in a Pol II dependent manner (Sigman *et al*., 2021). Presumably, the interaction of AGO4:siRNA:PolV transcript attracts de novo methyltransferase DRM2 in order to impose a first wave of cytosine methylation (Zhong *et al*., 2014). POL IV may be involved in the transcription of the methylated locus, generating short transcripts that are copied by RDR2 and processed by DCL3 into additional 24-nt siRNAs that amplify RdDM in a self-reinforcing loop (Zhai *et al*., 2015), resulting in the dense methylation of all cytosines in any sequence context: CG, CHG, CHH. Once de novo established at the presence of RNA triggers, CG and CHG may be mitotically/meiotically maintained even at the absence of the RNA triggers by the mere action of MET1 and CMT3, respectively (Aufsatz *et al*., 2004; Cao *et al*., 2003). However, CHH methylation cannot be maintained at the absence of RNA triggers, thus being the hallmark of de novo RdDM (Pelissier *et al*., 1999). Importantly, cytosine methylation attracts histone deacetylases and methyltransferases that lead to histone modifications of the methylated region, especially if it is a promoter region (Mirouze and Paszkowski, 2011; Zemach and Grafi, 2007). Thus, while DNA methylation in a coding region may not affect transcription, DNA methylation in a promoter region may ensue histone modifications and heterochromatinization, resulting in TGS (Wassenegger, 2005). However, while both transgenic and endogenous promoters are prone to RdDM, endogenous promoters are less prone to subsequent chromatin modifications and TGS for reasons that are not clear (Okano *et al*., 2008). Interestingly, the few endogenous promoters that are prone to TGS derive from tissue-specifically expressed genes (Eamens *et al*., 2008).

We have previously shown that high pressure spraying technology (HPST) is a very powerful tool to modify gene expression in plants by leading to PTGS of selected targets (Dalakouras *et al*., 2016; Uslu *et al*., 2021; Uslu and Wassenegger, 2020). Recently, the first example of spray-induced epigenetic modifications (SPIEM) has been reported, in which high pressure spraying of a 333 bp dsRNA triggered promoter methylation in *N. benthamiana* (Dalakouras and Ganopoulos, 2021). However, whether high pressure spraying of smaller RNAs may also trigger RdDM has not been reported, until this study. Here, we provide evidence that high pressure spraying of 24-nt siRNAs designed to target the CaMV 35S promoter triggered promoter RdDM and, importantly, TGS. Our approach of SPIEM by siRNAs rather than longer dsRNAs will most likely minimize epigenetic off-target effects in target and non-target organisms and as such, could well be employed in modern GMO-free, RNA-based, epi-breeding approaches (Dalakouras and Vlachostergios, 2021).

## RESULTS

We designed a 35S-promoter targeting 24nt-siRNA containing 2nt overhangs on both strands, mapping 21nt upstream of TATA-box between -114 and -89 positions with respect to the GFP transgene in 16C *Nicotiana benthamiana* line (Fig 1a). The 24nt-siRNA was sprayed at high pressure (4-6 bars) in 4 independent experiments on 24 plants and 68 leaves (Table 1). Only 8 plants with at least one silenced leaf and 14 leaves with at least two silenced spots were detected (Fig 1b). The silenced spots were mostly very tiny, scattered, and had orange color, which was remarkably different from the 22nt-RNA induced PTGS (Fig 1c) (Uslu et al 2021). In order to find out whether PTGS underlies the mild silencing, sRNA sequencing was performed to detect gene body transitivity. For this purpose, 6 leaves with the most silenced spots were selected among the 14 silenced leaves (Fig 1c). A rough rectangle containing as much silence spots as possible were cut out from the leaves by a razor. Silenced areas from two leaves were pooled for each of three biological replicates for sRNA sequencing.

**Table1.** 24nt-siRNA Spraying Scheme. In four independent HPST experiments, 8/24 plants showed silencing in at least one leaf per plant and 14/68 leaves showed at least two silencing spots. However, the efficiency of transgene silencing changes from one experiment to another.

| 24-nt siRNA HPST | Sprayed<br>Plant | Silenced<br>Plants | Sprayed<br>Leaves | Silenced<br>Leaves |
| --- | --- | --- | --- | --- |
| Experiment 1 | 12 | 4 | 36 | 6 |
| Experiment 2 | 4 | 1 | 12 | 3 |
| Experiment 3 | 4 | 0 | 8 | 0 |
| Experiment 4 | 4 | 3 | 12 | 5 |
| Total | 24 | 8 | 68 | 14 |

**Figure 1.**
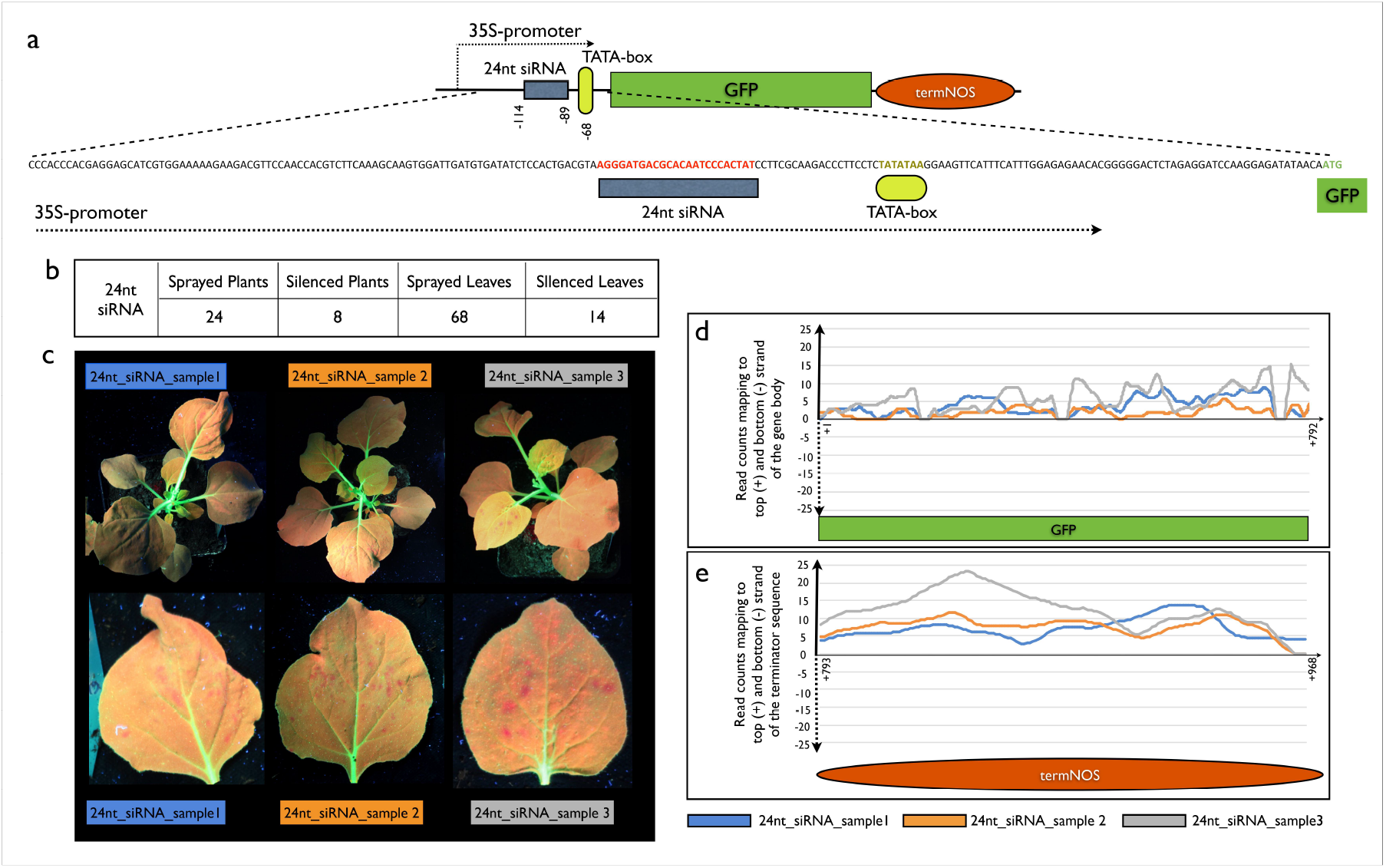
Transgene silencing by 24nt-siRNA HPST and sRNA signature: (a) The schematic representation of GFP locus in 16C plants shows that 24nt-siRNA is designed on the 3’ side of the 35S promoter. Below the 200nt of the 35S promoter sequence is given with the corresponding 24nt siRNA target site, TATAbox, and ATG translational start site of the GFP respectively(b) The summary of 4 independent experiments is indicated. At least two silenced areas are required to fulfill the silenced leaf criteria and at least one silenced leaf is counted as silenced plant (c) Three strongest examples of transgene silencing upon 24nt-siRNA delivery by HPST. The top panel shows the whole plants under UV light 6dps, and the bottom panel shows the silenced leaves, which are also used in sRNA sequencing and bisulfite analysis. (d) The line-chart indicates the total number of reads in a window of 10 nucleotides mapping to the top and the bottom strand of the GFP coding sequence in each biological replicate. We could not detect any reads mapping to the opposite strand, excluding any possible processing of the target GFP by RNAi. € The line-chart indicates the total number of reads in a window of 10 nucleotides mapping to the top and the bottom strand of the NOS terminator in each biological replicate. We could not detect any reads mapping to the opposite strand, excluding any possible processing of the target GFP and terminator sequence by RNAi.

sRNA reads mapping to the GFP coding sequence, and the terminator site showed that all the reads were in the same direction of the GFP transcript (Fig 1d,1e). These reads come from the degradation of the GFP and even detectable in water sprayed 16C plants (Uslu et al 2020). The lack of reads mapping to the negative strand suggested that the transitive silencing was not at a detectable level and could not support the hypothesis that PTGS underlies the silencing upon 24nt-siRNA spraying (Fig 1d, 1e). The sRNA reads mapping to the CaMV 35S promoter sense strand between -37 and - 1 are due to the transcription start site at the end of the 35S promoter (Fig 2a). On the other hand, the vast majority of the sRNA were mapped to the target site (Fig 2a). Since it is not possible to exclude the 24nt-sRNA on the leaf surface, we cannot identify the 24nt-siRNA, which was delivered into the cells. Nevertheless, approximately one percent of the reads mapping to the target site are longer than 24nt and expands to the neighboring sites (Fig 2b). Half of these reads are at least 30nt long and they are mapped on both positive and negative strand (Figure 2b). These RNAs, longer than 24nts are likely to be secondary RNAs produced by RdDM machinery upon HPST of 24nt siRNA.

**Figure 2.**
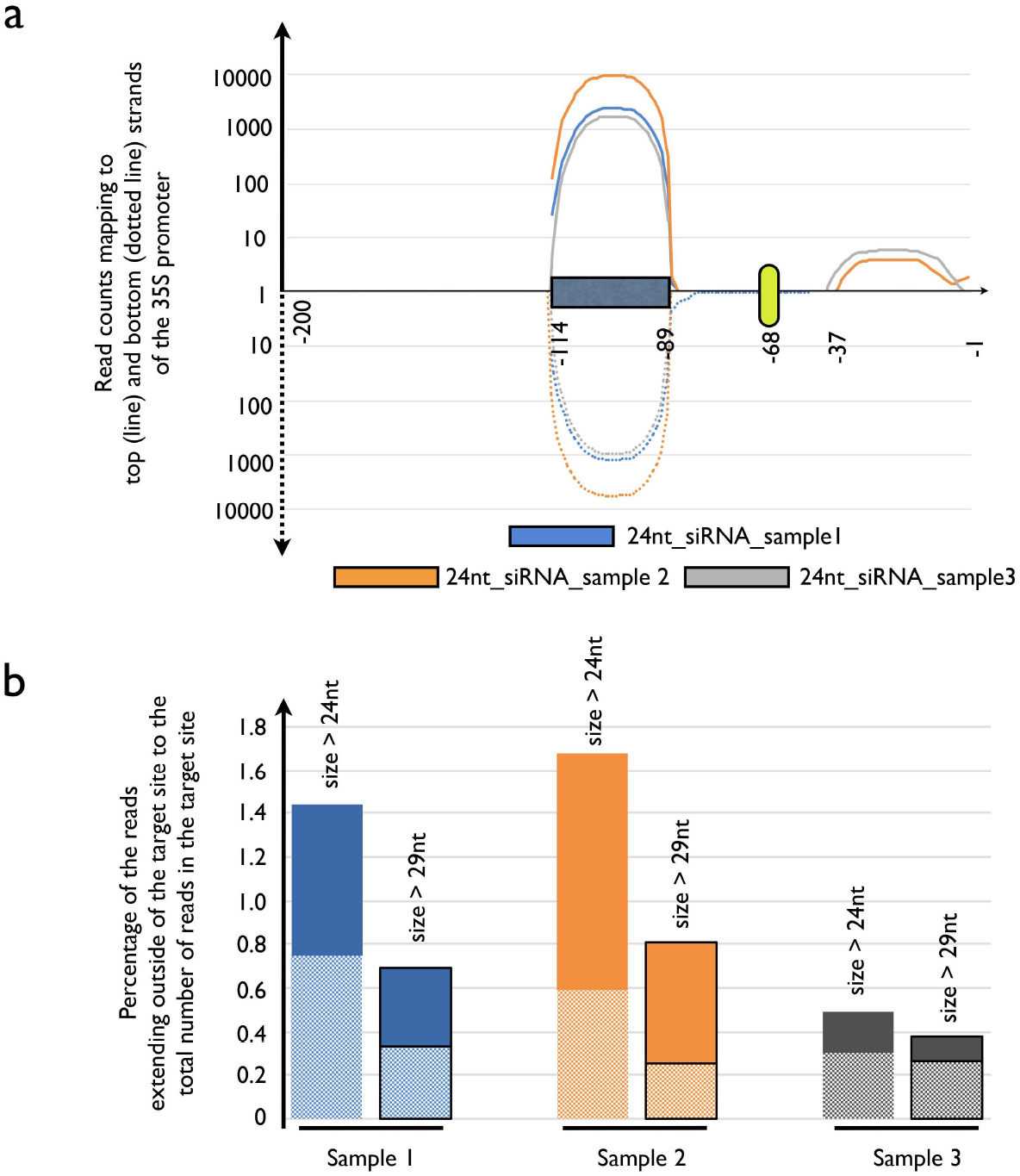
sRNA distribution in the CaMV 35S promoter upon 24nt siRNA spraying. (a) The total read counts mapping to the CaMV promoter is given in logarithmic scale. In the top panel, the reads mapping to the sense strand were shown with lines are shown for each biological replicate. In the bottom panel, the reads mapping to the anti-sense strand were shown with doted lines are shown for each biological replicate. In agreement with the previous literature 30nt downstream of the TATAbox is a transcriptional start site and at position -37 and -35, sense strand reads are shown (Xu et al 2016). (b) The secondary RNA reads, which are at least 25nt long and at least 30nt long are shown for each biological replicate. The solid colors on the top of the bars indicate the percent ratio of reads expanding outside the target site to the total reads mapping to the target site. The dotted patterns on the bottom of the bars indicate the percent ratio of reads expanding outside the target site to the total reads mapping to the target site.

In order to find out whether 24nt-siRNA could induce DNA methylation at the target site and in the vicinity of the target site, we performed bisulfite sequencing on the top and the bottom strand of the 35S promoter containing the target site. Approximately 50mg of leaf material was collected around the silenced leaf areas. As a control water sprayed and 22nt-siRNA164, which target GFP gene body, were used. For analysis 35S promoter was divided into three parts: The target site (T-site), which matches the 24nt-siRNA, the neighboring sites (N-sites), which contains five cytosine residues on both sides of the T-site, and thirdly the distal areas (Off-site), which lie outside of the N-sites (Fig 3a, 3b).

**Figure 3.**
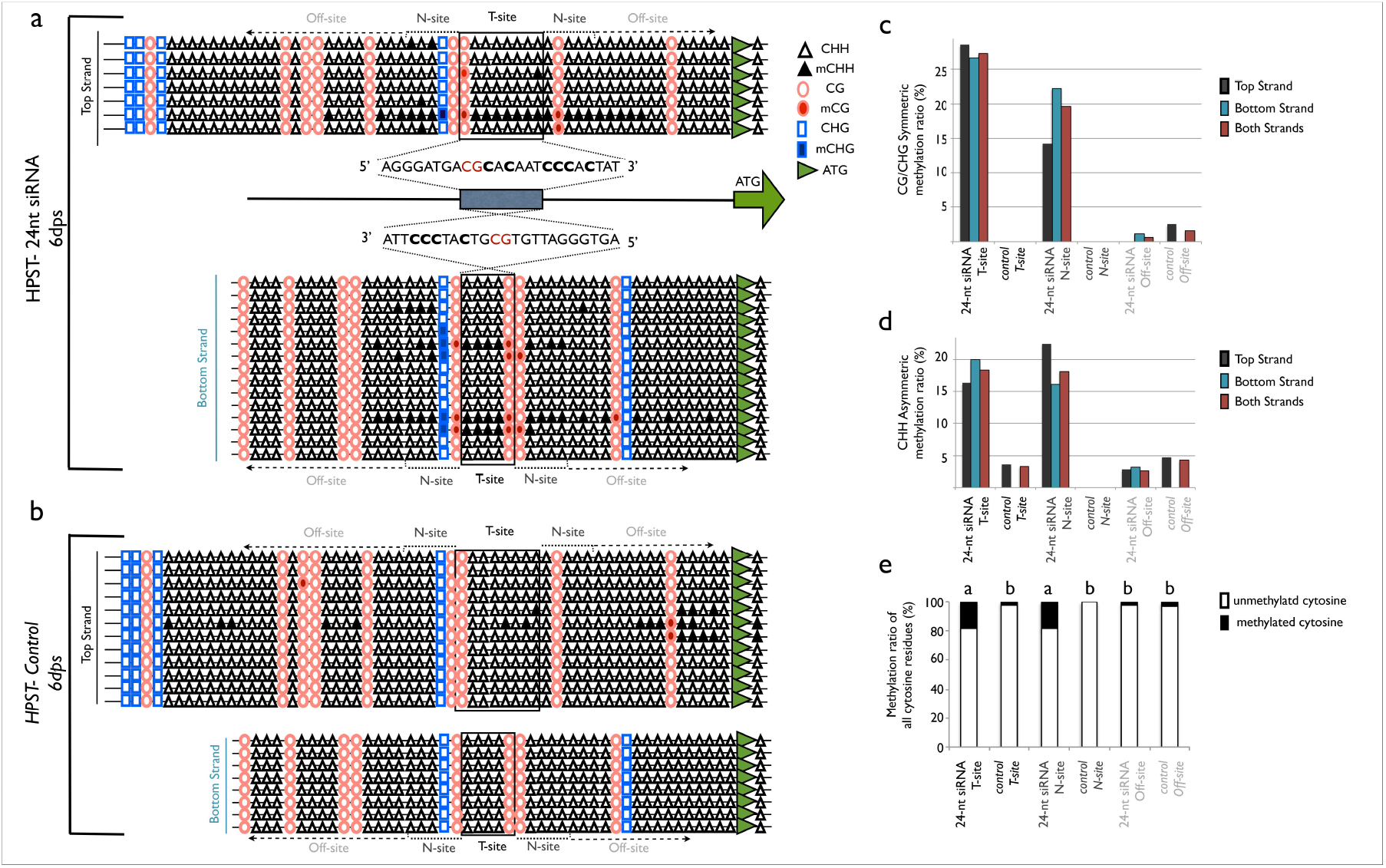
35S promoter methylation analysis upon SPIEM by 24nt-siRNA: Bisulfite sequencing analysis is schematized by triangle, circle, and rectangles representing asymmetric CHH and symmetric CG and CHG sequences. The filled shapes indicate non-converted cytosines upon bisulfite treatment, which are the methylated residues. The green arrow shows the ATG translation start site of GFP transgene in 16C plants. In the bisulfite sequences, Target sites (T-sites), marked by black rectangle, show where the 24nt-siRNA map. Neighboring N-sites expand five cytosines on both sides of the T-site, and Off-sites are any cytosines outside of the T-and the N-sites. The Off-sites shown in the graph are trimmed for visualization reasons. Multiple identical sequences from the same biological replicates are represented by one single read - unless indicated (a) Methylation upon HPST of 24nt-siRNA is shown. In the top panel, methylation of the Top Strand is given. 7 sequences originate from 3 biological replicates and minimum 2 technical replicates. In the middle panel, 24nt-siRNA sequence is given and the symmetric CG sequence, present in both top and bottom strands is shown in red. In the lower panel bottom strand methylation (3’ to 5’) is given 15 sequences originate from 4 biological replicates and two technical replicates are shown. (b) Methylation in non-treated and upon HPST of water is shown. In the top panel, methylation of the top 35S-promoter strand is given. 12 sequences originate from 2 biological replicates (non-treated) with minimum 2 technical replicates and 3 biological replicates (water sprayed) with minimum 2 technical replicates shown. In the lower panel bottom strand methylation is given 8 sequences originate from 5 biological replicates. Since none of the sequences showed methylation, we did not eliminate any of the reads. (c) The symmetric methylation ratio is calculated for 24nt-siRNA treated and control samples in T-site, N-site, and Off-site for both strands and in cumulatively. (d) The asymmetric methylation ratio is calculated for 24nt-siRNA treated and control samples in T-site, N-site, and Off-site for both strands and in cumulatively. (e) Cumulative cytosines are calculated in T-site, N-site, and Off-site and pairwise statistical comparison has been done by Fisher’s Exact Test based on the number of methylated and non-methylated cytosines with a significance level of p<0.01.

Bisulfite sequencing showed that one fourth of the symmetric cytosine residues (CG and CHG) are methylated at the target site, regardless of the strand direction, whereas no CG or CHG methylation at of the target site was detectable in the controls (Figure 3c). Besides, approximately one fifth of the symmetric cytosine residues are methylated in the N-sites, while the N-sites were completely unmethylated in the controls. The symmetric methylation in the Off-site was less than 3 percent for both 24nt-siRNA treated and control samples.

In terms of asymmetric methylation, we detected approximately one sixth of the CHH residues were methylated at the T-site and N-site upon 24nt-siRNA spraying. However, control group showed less than 5 percent CHH methylation in the T-site and no methylation at the N-site. Off-site CHH methylation was not significantly different in both 24nt-siRNA treatment and controls. (Fig 3d).

When all the cytosine residues are taken into consideration, one fifth of the cytosine residues are methylated upon treatment in T-site and N-site, which is significantly higher than the control methylation sites (Fisher’s exact test: p<0.01). On the other hand, the off-site methylation is not significantly different upon spraying 24nt-siRNA and controls (Figure 3e)

In order to increase the efficiency of 24nt siRNA delivery, we used carbon dots, which was shown to deliver 21 and 22nt siRNAs into plant cells with high efficiency (Schwartz et al 2020, Hendrix et al 2021). We have observed that low pressure spraying of 22nt-siRNAs in combination with carbon dots produce wider silenced areas when compared to the 22nt-siRNAs delivered by HPST (Fig 4a). High pressure spraying with carbon dots lead to too strong wounding and necrosis in the leaves to deduce its silencing efficiency (data not shown). Therefore, we investigated whether carbon dots could increase the delivery of 24nt-siRNAs upon low pressure (LP) spraying. 24-nt siRNAs delivered with carbon dots (CD) did not lead to any silencing phenotype (Fig 4a). Moreover, we also could not detect any T-site and N-site methylation upon LP-CD spraying, indicating that low pressure spraying even in combination with carbon dots could not induce RdDM via exoRNAs (Fig 4b).

**Figure 4.**
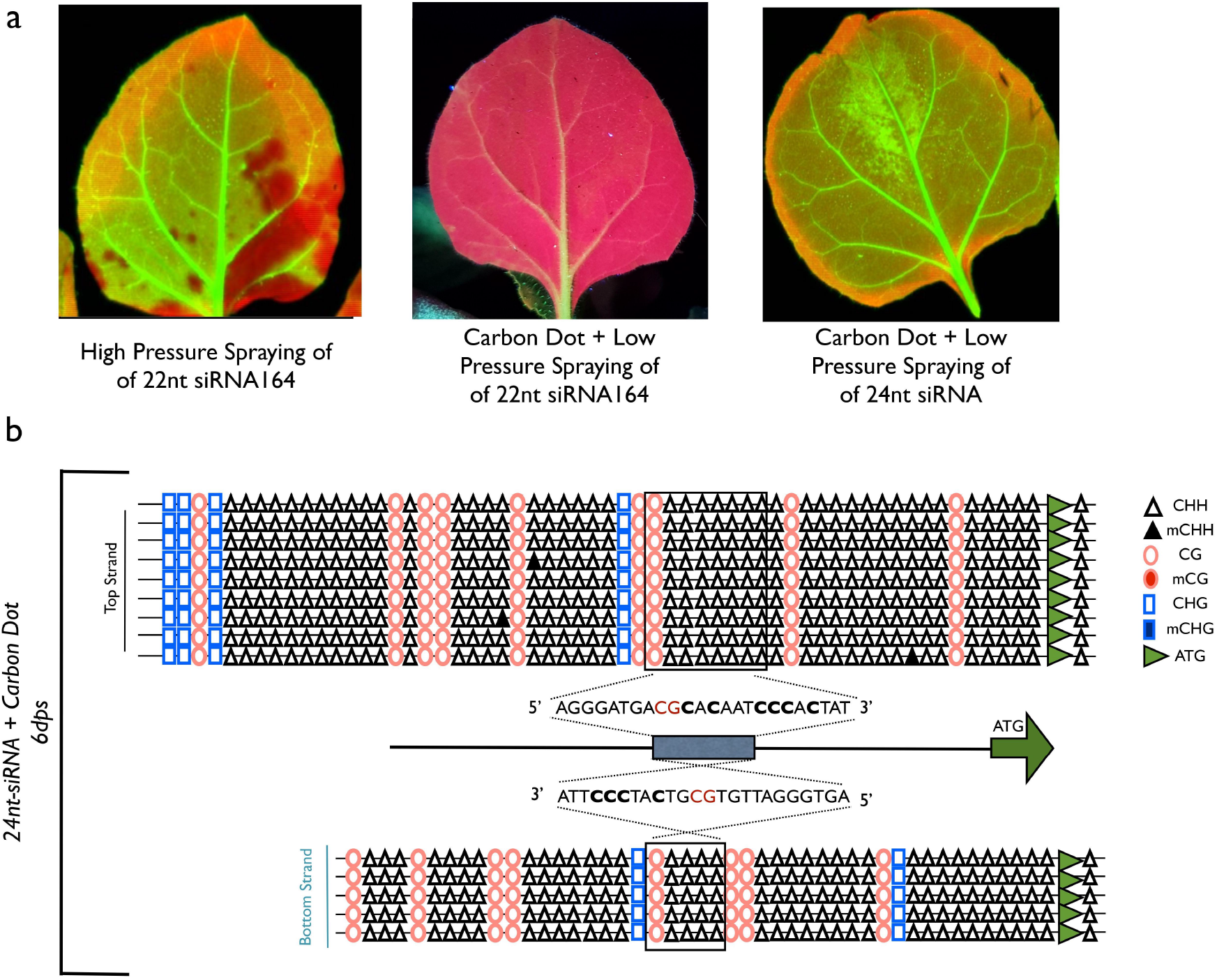
Silencing and 35S promoter methylation analysis upon siRNA-Carbon Dot complexes. (a) The silencing phenotypes of 16C plants are shown. 22nt-siRNA164 by HPST (Uslu VV et al 2021) leads to consistent local silencing covering approximately 20% to 80% of the leaf surface. 22nt-siRNA164 by Carbon Dot and Low-Pressure Spraying (CD-LP) can lead to local silencing of the whole leaf area. However, no silencing upon 24nt-siRNA by CD-LP is observed in three independent experiments. (b) The top and the bottom panel shows the methylation pattern of the 35S promoter in the top and the opposite strand, respectively. The target site is shown in black rectangle. 3 biological replicates of 24nt-siRNA by CD-LP are shown with minimum 2 technical replicates. In the bottom panel, non-treated, LP with only water, only Silwett 77, Carbon Dot, 22nt-siRNA by CD-LP is given. Since no methylation was observed in the bottom strand, one example of each treatment is shown in the control.

## DISCUSSION

A quarter of the leaves, sprayed with 24nt-siRNA, showed mild to very mild silencing phenotype. The lack of secondary siRNAs in the gene body and the terminator but the presence of secondary RNAs in the promoter suggests that TGS but not the PTGS underlies the mild silencing of the leaves upon 24nt-siRNA spraying. Although, it is still unexpected to have visible silencing via TGS as early as 6dps based on previous work (Dalakouras and Gannopoulos 2021), based on the half-life of the GFP mRNA and GFP protein, theoretically it is possible to detect silencing at 6dps (Sacchetti et al 2001, Corish and Tyler-Smith 1999). Therefore, the orange rather than the red tone of the silenced areas under UV light may be due to incomplete silencing or leaky expression in the cells, where 35S promoter is partially methylated by the 24nt siRNA.

Previously published data on inducing promoter methylation via HIGS, VIGS, and exogenous dsRNA application show conflicting results on the border of the methylated area: While, in most of the cases, the methylation site expands the target site (Dalakouras et al 2016, Zilberman et al 2004, Vaistij et al 2002), dsRNA delivery and intronic hairpin RNA induced RdDM did not show any indication of spreading of methylation outside of the target site (Dalakouras and Gannopoulos 2021, Dalakouras et al 2009, Dalakouras 2010, Vogt et al 2004). Our data on HPST of 24nt siRNAs shows that the methylation can spread at least up to five cytosines residues neighboring the target site on both 5’ and 3’ sides in both strands. Moreover, we also detect secondary small RNAs extending over the neighboring areas (Fig 2b), suggesting that a single 24nt siRNA population can trigger production of secondary siRNAs in the target site by RdDM machinery. Considering that the secondary RNAs has at least 22nt match with the target site, the synthesis of the secondary RNAs is primed by the initial 24nt trigger siRNA.

CG and CHG methylation are maintained after cell division primarily by MET1, and CMT3, respectively, while the CHH methylation is a de novo process, which requires DRM2 and CMT2 activity and the trigger to be present (Erdmann and Picard 2020). In our bisulfite experiments, the CG and CHG methylation is slightly higher than the CHH methylation (Figure 3d,3e). This suggests that either the cell division is slow so CHH methylation is largely maintained or the methylation took place shortly before the sample collection. Considering that the mature leave cells divide slowly and there is a mild silencing phenotype, the slight difference between CG/CHG methylation and CHH can be interpreted as a result of slow cell division.

It has been shown that the dsRNA spraying did not induce any methylation at 3dps (Dalakouras and Gannopoulos 2021). However, at 10dps, 40% of the samples were heavily methylated at the target site. We showed that 24nt-siRNA spraying lead to target site heavy methylation of the 28% percent of the samples at 6dps, whereas 72% of the samples were not methylated at all (Fig 3). Our findings suggest that whenever the delivery of the siRNA into the nucleus took place, it could trigger methylation machinery at high efficiency. However, the siRNAs were taken up by only a fraction of cells. Therefore, the T-site and N-site methylation pattern across different samples appears as a bimodal distribution of heavy silencing and no silencing at all, as observed previously for exo-dsRNA delivery (Dalakouras and Gannopoulos 2021).

An important difference between PTGS and TGS is the cellular compartments that these mechanisms take place. PTGS is executed mostly in the cytosol, but the nucleus is the critical compartment for TGS. Carbon Dots (CD) increase the efficiency of PTGS in local leaves upon 22nt siRNA spraying when compared to high pressure spraying (Fig 4a). However, we did not detect any methylation upon low pressure spraying of the 24nt siRNA with CDs (Fig 4b). CDs are effective RNA carriers into the cytosol but not into the nucleus. It has been hypothesized that siRNA-CD complexes are carried into the cytosol by endocytosis and due to the “proton-sponge” effect, siRNAs can be released to the cytosol by osmotic swelling (Schwartz et al 2020), possibly restricting the delivery to the nucleus. On the other hand, the delivery of 24nt-siRNA into nucleus can effectively be achieved upon high pressure spraying most likely via mechanical penetration of the siRNA at 4-6 bar pressure.

In this study we introduced the proof of principle for using 24nt siRNAs for SPIEM at the target sites. We showed that we could target 20% of the leaf cells by high pressure spraying. Nevertheless, there is room for improvement for enhancing the efficiency of RNA delivery by novel RNA carrier molecules alone or in combination with HPST. Further work is needed to elucidate repressive histone modifications at the sites *de novo* methylated by 24nt siRNA. In contrast to 24nt siRNA or dsRNAs, SPIEM by trans-acting non-coding RNAs have the potential to activate target gene expression. Therefore, the spray induced epigenetic modification (SPIEM) method shown in this manuscript could potentially be used in epigenetic-based plant breeding approaches practically for any crop plant. However, to be able to translate this technology into application the RNA delivery and the methylation efficiency at the meristem tissue and the germline developed from the targeted meristem should be investigated in the future.

## MATERIALS and METHODS

### Plant Growth and Phenotyping

Nicotiana benthamiana 16C plants were grown in 25°C, 16h light - 8h dark cycle. The spraying experiments were performed between 20- and 25-days post germination, which correspond to 4- to 6-true leaf stage. The silencing phenotype at 6dps was visualized under UV light (Blak-Ray B-100 AP Lamp, www.uvp.com) and images are obtained by Canon EOS700D (18-55mm) in Aperture Priority Mode (A:10) or Molecular Imager® PharosFX™ Systems (www.bio-rad.com) with GFP and Cy5 fluorescence. The images were overlayed by FIJI (www.imagej.net)

### 22nt- and 24-nt siRNA Spray

4-6 bar pressure (depending on the leaf size) and 0.5-1bar pressure provided by METABO Elektra Beckum Classic 250 compressor (www.metabo.com) is used for HPST and Low-Pressure (LP) Spray, respectively. Airbrush pistol (CONRAD AFC-250A and CONRAD HP-200) is used to deliver the siRNAs from a distance of than 1cm. 24-nt siRNA was provided by Metabion, 22nt-siRNA is the siRNA164 in Uslu VV et al 2021. The RNA was dissolved in water for naked RNA spraying at high pressure and Silwett L77 + CD Spray buffer (10mM MES, 20mM glycerol, pH:5.7) is used for CD spray.

### RNA extraction and small RNA Sequencing

At 6dps, silenced areas were excised out from 2 leaves, showing mild silencing and were pooled per biological replicate. Three biological replicates of approximately 50mg leaf material were crashed using metal beads and small RNAs were extracted using mirVana miRNA extraction kit according to manufacturer’s instructions (www.thermofisher.com). sRNA-seq and bioinformatic analysis were performed by GenXPro, Frankfurt (http://genexpro.net) as indicated in the literature (Uslu VV et al 2021).

### Bisulfite Sequencing

Genomic DNA is extracted by PureLink™ Genomic Plant DNA Purification Kit (www.thermofisher.com, not available since June, 2021) and Plant DNA Preparation-Solution Kit (www.jenabioscience.com from June, 2021 on).

Zymo EZ DNA Methylation Direct Kit (www.zymoresearch.de) is used to convert unmethylated cytosines. Conversion reaction was performed for 16h at 65°C. 35S promoter PCR template is converted by bisulfite kit to evaluate conversion efficiency (>99%). DreamTaq DNA polymerase (www.thermofisher.com) is used to amplify the bottom and top strand with the following primer pairs, respectively: 1995_24nt_35Sp_oppStr-Fwd: 5’-TCCCAAARATRRACCCCCACCCAC-3’, 1995_24nt_35Sp_oppStr-Rev: 5’-TGAAAAGATGAGAAAGAGAAAAAGATTAG-3’, 2103_24nt_35S-TopStr-Fwd2: 5’-GAAGATAGTGGAAAAGGAAGGTG-3’, and 2105_24nt_35S_TopStr-Rev2: 5’-AAAARTTCTTCTCCTTTACT-3’ The PCR products after 35 cycles are cloned into sequencing vector by CloneJET PCR Cloning Kit (www.thermofisher.com). T7 primer is used for sequencing the insert. Initial analysis of methylation has been performed by CyMATE (www.cymate.org) and the data is further curated to exclude sequencing errors manually.

## Conflict of Interest

The authors declare that the research was conducted in the absence of any commercial or financial relationships that could be construed as a potential conflict of interest.

## Author Contributions

GK, MWass, and VVU conceived and designed the experiment. MW, LH, BE, and VVU performed experiments. LH performed the CD synthesis and CD spraying. MW and VVU performed the data analysis and VVU wrote the manuscript.

## Funding

The project is funded by Federal Ministry of Education and Research (BMBF) Grant No: 03180531

## Acknowledgments

We would like to thank our colleagues Alexandra Baßler, Michele Wassenegger, Saeideh Ebrahimi at the Bioökonomie/Epigenetic Department of AgroScience for valuable input and discussion. We would like to acknowledge Benedikt Eisenger for his help in setting up and testing the bisulfite assay. In addition, we would like to thank Athanasios Dalakouras for critical reading of the manuscript and fruitful discussions.

## Data Availability Statement

The sRNA-seq data currently available upon request and will be available in the GEO database.

## Notes

### Competing Interest Statement

The authors have declared no competing interest.

